# A theoretical justification for single molecule peptide sequencing

**DOI:** 10.1101/010587

**Authors:** Jagannath Swaminathan, Alexander A. Boulgakov, Edward M. Marcotte

**Author notes:** These authors contributed equally (Jagannath Swaminathan, Alexander A. Boulgakov).

## Abstract

The proteomes of cells, tissues, and organisms reflect active cellular processes and change continuously in response to intracellular and extracellular cues. Deep, quantitative profiling of the proteome, especially if combined with mRNA and metabolite measurements, should provide an unprecedented view of cell state, better revealing functions and interactions of cell components. Molecular diagnostics and biomarker should benefit particularly from the accurate quantification of proteomes, since complex diseases like cancer change protein abundances and modifications. Currently, shotgun mass spectrometry is the primary technology for high-throughput protein identification and quantification; while powerful, it lacks high sensitivity and coverage. We draw parallels with next-generation DNA sequencing and propose a strategy, termed fluorosequencing, for sequencing peptides in a complex protein sample at the level of single molecules. In the proposed approach, millions of individual fluorescently labeled peptides are visualized in parallel, monitoring changing patterns of fluorescence intensity as N-terminal amino acids are sequentially removed, and using the resulting fluorescence signatures (fluorosequences) to uniquely identify individual peptides. We introduce a theoretical foundation for fluorosequencing, and by using Monte Carlo computer simulations, we explore its feasibility, anticipate the most likely experimental errors, quantify their potential impact, and discuss the broad potential utility offered by a high-throughput peptide sequencing technology.

**AUTHOR SUMMARY:** The development of next-generation DNA and RNA sequencing methods has transformed biology, with current platforms generating >1 billion sequencing reads per run. Unfortunately, no method of similar scale and throughput exists to identify and quantify specific proteins in complex mixtures, representing a critical bottleneck in many biochemical and molecular diagnostic assays. What is urgently needed is a massively parallel method, akin to next-gen DNA sequencing, for identifying and quantifying peptides or proteins in a sample. In principle, single-molecule peptide sequencing could achieve this goal, allowing billions of distinct peptides to be sequenced in parallel and thereby identifying proteins composing the sample and digitally quantifying them by direct counting of peptides. Here, we discuss theoretical considerations of single molecule peptide sequencing, suggest one possible experimental strategy, and, using computer simulations, characterize the potential utility and unusual properties of this future proteomics technology.

## INTRODUCTION

The basis of “next-gen” DNA sequencing is the sequencing of large numbers of short reads (typically 35–500 nucleotides) in parallel. Currently available next-generation sequencing platforms from Pacific Biosciences [1] and Helicos [2] monitor the sequencing of single DNA molecules using fluorescence microscopy and can allow for approx. one billion sequencing reads per run (e.g., for Helicos). Unfortunately, no method of similar scale and throughput exists to identify and quantify specific proteins in complex mixtures, representing a critical bottleneck in many biochemical, molecular diagnostic, and biomarker discovery assays. For example, consider the case of cancer biomarker discovery: nucleic acid mutations underlie nearly all cancers. However, these variants are embodied by proteins and are often expressed in bodily compartments (saliva, blood, urine) accessible without invasive biopsies. The use of protein biomarkers to diagnose, characterize, and monitor most, if not all, cancers [3] would be significantly advanced by an approach to sensitively identify and quantify proteins in these compartments. Indeed, the value of diagnostic biomarkers is clearly seen in the utility of detecting thyroglobulin for monitoring thyroid cancer, and in administering Herceptin specifically for breast cancers overexpressing HER2/neu [4]. Techniques applied to this problem, including mass spectrometry (MS) and antibody arrays, often lack sufficient sensitivity and intrinsic digital quantification to be effective [5]. What is urgently needed is a massively parallel method, akin to next-gen DNA sequencing, for identifying and quantifying individual peptides or proteins in a sample.

Unlike the polymerase chain reaction (PCR) for nucleic acids, no methods exist for template-directed amplification of proteins. Hence, advances in proteome analysis sensitivity and throughput often focus on enhancing detectors, or even on inferring proteomes based on measuring translating mRNAs [6]. The most sensitive current methods for discriminating specific arbitrary protein sequences, such as mass spectrometry, typically exhibit attomole or femtomole sensitivities [7], although new detection modalities for detecting specific single molecule sequences are being developed [8]. Nonetheless, current proteomics technologies tend to lag the analogous nucleic acid sequencing technologies by 3 to 5 orders of magnitude in both throughput and sensitivity. Recently, researchers have attempted to identify single protein molecules using nanopores [9, 10], although the challenge of deconvoluting electrical signals to sequence peptides is currently unsolved.

In principle, extending classic Edman degradation protein sequencing methodology to the single-molecule level could potentially allow billions of distinct peptides to be sequenced in parallel, thereby identifying proteins composing the sample and digitally quantifying them by direct counting of peptides. Edman degradation, first described by Pehr Edman in 1949 [11], is a standard method to determine the amino acid sequence of a purified peptide, in which the amino-terminal (N-terminal) amino acid residue is labeled and cleaved from the peptide without disrupting the peptide bonds between other amino acid residues [12]. In the conventional approach, the freed amino acid is identified (e.g., by chromatography), and successive cycles of this procedure reveal the sequence of amino acids in the peptide.

In this paper, we have tried to anticipate how one might implement a single-molecule sequencing technology based on Edman degradation. We suggest one practical approach that would in principle be capable of generating partial peptide sequences in a highly parallel fashion, scalable to entire proteomes. Beyond biomarker discovery, such an approach would have broad applications across biology and medicine and could be as fundamental for proteins as, for example, PCR is for nucleic acid research. From a theoretical perspective, we discuss the many interesting features that data generated by such an approach would have, along with how such data might be interpreted and how sensitive the process might be to potential errors, which we model using Monte Carlo simulations.

## RESULTS AND DISCUSSION

### One possible strategy for implementing single molecule peptide sequencing

Figure 1 illustrates a proposed scheme for single molecule peptide sequencing. The key idea is to selectively fluorescently label amino acids on immobilized peptides, followed by successive cycles of removing peptides’ N-terminal residues (by Edman degradation) and imaging the corresponding decreases of fluorescence intensity for individual peptide molecules. The resulting stair-step patterns fluorescence decreases will often be sufficiently reflective of their sequences to allow unique identification of the peptides by comparison to a reference proteome.

**Figure 1:**
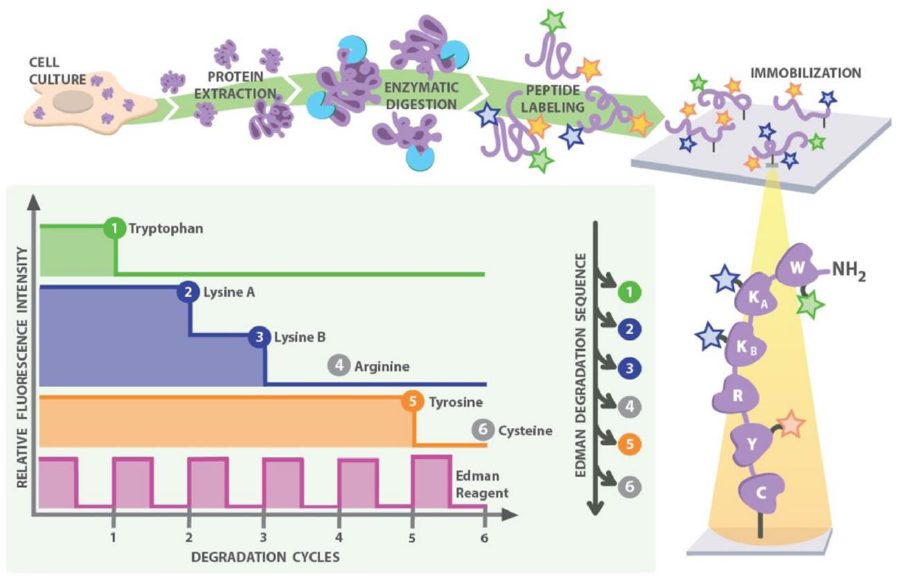
A strategy for single-molecule peptide sequencing. Proteins are extracted and digested into peptides by a sequence-specific endo-peptidase. All occurrences of particular amino acids are selectively labeled by fluorescent dyes (e.g., yellow for tyrosine, green for tryptophan, and blue for lysine residues), and the peptides are surface immobilized for single-molecule imaging (e.g. by anchoring *via* cysteine). The peptides are subjected to cycles of Edman degradation; in each cycle, a fluorescent Edman reagent (pink trace) couples to and removes the most N-terminal amino acid. The step drop of fluorescent intensity indicates when labeled amino acids are removed, which in combination with the Edman cycle completion signal, gives the resulting *fluorosequence* (e.g., “WKKxY…”). Matching this partial sequence to a reference protein database identifies the peptide.

In more detail, proteins in a complex mixture are first proteolytically digested into peptides using an endo-peptidase of known cleavage specificity. Select amino acid types (e.g. lysine, tryptophan or tyrosine) are covalently labeled with spectrally distinguishable fluorophores, each being specific (by reactivity) to the given amino acid side chain. Labeled peptides are immobilized on a glass surface, as for example *via* the formation of a stable thioether linkage between a maleimide functionalized surface and the thiol group on cysteine residues [13]. The choice of peptidase, labeled amino acids, and anchor all convey information about the identity of a peptide and thus can be optimized for maximum effect. Using techniques such as Total Internal Reflection Fluorescence (TIRF) microscopy, individual peptide molecules can be imaged on such a surface, and the fluorescence intensity across all fluorophore channels can be determined for each peptide on a molecule-by-molecule basis. By monitoring decreases in fluorescence intensity following cycles of Edman degradation, we can determine the relative positions of labeled amino acids in the peptides, and thereby obtain a partial peptide sequence. This scheme might be improved by using a fluorescent Edman reagent whose coupling and decoupling can be observed, enabling the successful completion of each Edman cycle to be monitored for every single peptide, providing an additional error check.

We term the pairing of an Edman degradation cycle and the subsequent observation for changes in fluorescence an *experimental cycle* (see **Box 1** for definitions of terminology). The observed sequence of luminosity drops in fluorescence across experimental cycles is a *fluorosequence*; the technique itself is thus *fluorosequencing*. For the example shown in Figure 1, the fluorosequence is “WKKxY”. Mapping the partial sequence back to a *reference proteome* of potential proteins, such as might be derived from a genome sequence, would determine if the fluorosequence uniquely identifies a peptide, and ultimately, its parent protein.

Commercially available TIRF microscopes can easily monitor fluorescence changes for millions of individual peptide molecules [14] and are not dissimilar to early variants of next-generation DNA sequencers [2]. By increasing peptide density and acquiring TIRF images over a large surface area, one could in principle obtain fluorosequences for millions or billions of peptides in parallel. Critically, this approach would be intrinsically quantitative and digital, based on counting repeat peptide observations, in much the same way NextGen RNA sequencing is for identifying and quantifying RNA transcripts.

### Under ideal conditions, even partial amino acid sequences are informative

Computer simulations of variations of this scheme confirm that fluorosequences can be quite information rich; even relatively simple labeling schemes, employing only 1 to 4 amino acid-specific fluorescent labels, can yield patterns capable of uniquely identifying at least one peptide from most of the known human proteins (Figure 2). For these simulations, we considered only labeling schemes based on known differences in side-chain reactivity and available amino acid-specific targeting chemistry [15], such as the reactivity of diazonium groups for tyrosines [16].

**Figure 2:**
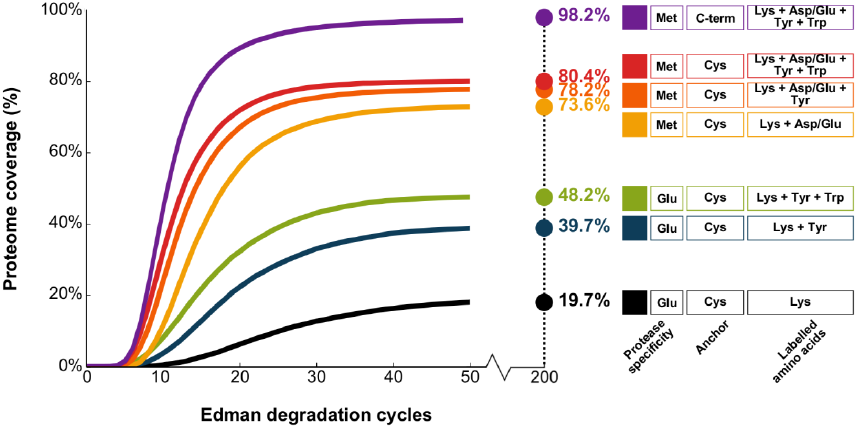
Simulations of ideal experimental conditions suggest relatively simple labeling schemes are sufficient to identify most proteins in the human proteome. Each curve summarizes the fraction of human proteins uniquely identified by at least one peptide as a function of the number of sequential experimental cycles (a paired Edman degradation reaction and TIRF observation). Here, we consider peptides generated by different proteases (*e.g.* Glu represents cleavage C-terminal to glutamic acid residues by GluC, Met represents cleavage after methionine residues by cyanogen bromide) and under different labeling schemes (*e.g.* Lys + Tyr indicates Lys and Tyr selectively labeled with two distinguishable fluorophores. Asp/Glu indicates both residues are labeled with identical fluorophores). Peptides are immobilized as indicated, with Cys representing anchoring by cysteines (thus, only cysteine-containing peptides are sequenced) and C-term representing anchoring by C-terminal amino acids. Increasing the number of distinct label types improves identification up to 80% within only 20 experimental cycles even when only Cys-containing peptides are sequenced; near total proteome coverage is theoretically achievable when cyanogen bromide generated peptides are anchored by their C-termini and labeled by a combination of four different fluorophores. Cycle numbers denote upper bounds, since each fluorosequence is not allowed to proceed past the anchoring residue (cysteine or C-terminus). Note also that the peptide length distributions change depending on the enzyme used for cleavage, with median lengths of 26 amino acids for cyanogen bromide, 8 for GluC and 10 for trypsin digests.

The above labeling schemes (anchoring peptides *via* internal cysteine residues) fail to achieve 100% coverage of the template proteome even after many experimental cycles under ideal conditions. The reason is two-fold: (a) Edman reactions cannot continue past the cysteine anchor or (b) the proteome contains paralogs and protein families differing at unlabeled amino acids that are hence indistinguishable. When simulations were repeated for the case of anchoring all cyanogen bromide cleaved peptides, not just cysteine-containing ones, by their C-termini, the coverage of the four-label scheme rose from 80% to 98% of the proteome (Figure 2, **top curve**). Moreover, when simulations were performed for the case of no proteolysis and anchoring each full length protein at its C-terminus, four of the tested multiple-label schemes (including schemes with only 2 label types) achieved over 96% coverage of the proteome within 200 experimental cycles. The remaining proteins were unidentified due to protein families being indistinguishable by the labeling schemes employed. These simulations thus confirm that single molecule fluorosequencing is intrinsically capable of identifying a majority of proteins in a proteome even when the number of label types is small.

It is also worth considering whether the linear scaling and dynamic range of photon detection by existing cameras might place a limit on the ability to discriminate luminosity drops in fluorescent intensity per peptide. For example, while it might be easy to discriminate a reduction from 5 to 4 fluorophores on a peptide, discriminating a reduction from 25 to 24 fluorophores could be difficult. However, the median count of labelable amino acids per peptide is often small. For example, when considering peptides generated by the protease GluC, this count ranges from approximately 2 (for lysines) to 7 (for glutamic acid/aspartic acid residues, which we assume are indistinguishable by reactivity for labeling purposes) (Figure 3). This range is well within the capacity of most modern cameras, since, in practice, TIRF microscopes equipped with CCD camera variants can count up to at least 13 fluorophores; that is, up to at least 13 copies of a given fluorophore per single molecule can be quantitatively distinguished [17]. Thus, peptides from typical proteomes should not be problematic in this regard.

**Figure 3:**
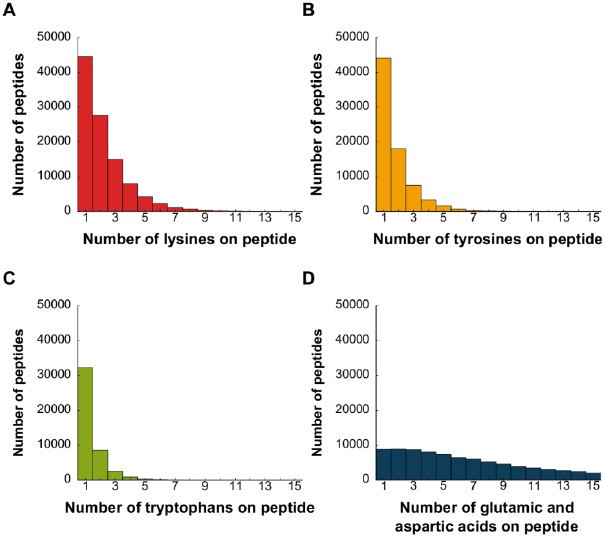
Typical proteolytic peptides have counts of labelable amino acids sufficiently low to sequence. Frequency histograms of amino acids in *in silico* proteolytic peptides for lysine **(A)**, tyrosine **(B)**, tryptophan **(C),** and glutamic acid/ aspartic acid **(D)** indicate low median values. Peptide sequences in A-C were generated *in silico* from the human proteome by GluC digestion, and those in D by cyanogen bromide digestion. Low counts of labelable amino acids per peptide are expected to increase the ability to discriminate removal of one fluorophore amongst many on a peptide.

### Anticipating the inevitable failures of dyes and Edman chemistry

Being a physico-chemical process, we can anticipate some of the most likely potential sources of error for an experimental implementation of the scheme. With errors, an observed fluorosequence would not reflect the true sequence of fluorescently labeled amino acids. Three of the most probable error sources are as follows:

a. **Failure of fluorophore attachment or emission causing apparent substitutions:** Steric constraints of peptides or reaction kinetics of fluorophore labeling chemistry might result in specific amino acid(s) not being covalently labeled. This scenario is equivalent to correctly coupled but non-emitting fluorophores, such as those observed in defective fluorophores [18]. In both circumstances, the position of a labelable amino acid would be misinterpreted as containing a non-labelable amino acid, *e.g.* the peptide “GK*EGK*” (where K* represents a labeled lysine) would mistakenly yield a fluorosequence “xxxxK” instead of “xKxxK”, for the dye failure at the first lysine.
b. **Photobleaching of labeled fluorophores causing apparent coupled double substitutions (“residue swaps”)**: The permanent photochemical destruction of dyes could also complicate the analysis. In this scenario, a labeled residue at one position is misinterpreted as an unlabeled residue because the label is lost by photobleaching, while another residue upstream in the peptide (typically unlabeled) is misinterpreted as being labeled because the photobleaching fluorophore loss coincides with that particular experimental cycle. This would shift the apparent position of the label upstream in the fluorosequence. For example, peptide GK*EGK* might be observed as xKKxx when the dye on the lysine at the fifth position photobleaches during the third imaging cycle. This situation reduces the ability to (i) reliably count the number of fluors lost during an experimental cycle, (ii) distinguish whether a change in luminosity results from fluorophore loss due to a genuine Edman degradation step or photobleaching, and (iii) identify which downstream fluorophore was extinguished if the loss is indeed due to photobleaching. Although fluorophore half-lives can be extended by use of oxygen scavenging systems [19], synthesis of stable dyes [20] or even surface modification [21], photobleaching is still a stochastic process and accounting for loss of fluorophores erroneously coincident with upstream Edman degradations would be critical to identification. Currently, there are many photo-stable dyes on the market. A recent study on the effects on dyes by oxygen radicals found that the half-life of Atto647 was roughly 3 minutes (corresponding to 180 experimental cycles at 1 second/cycle exposure) [22], while Atto655 showed a mean photobleaching lifetime of 8-20 minutes [23], corresponding to many hundreds of experimental cycles.
c. **Inefficiency of Edman degradation chemistry causing apparent insertions**: Optimization of Edman degradation over the past sixty years has resulted in efficiencies of >95% [24]. Nonetheless, failed cycles are expected at some non-zero rate and would yield an observation corresponding to no fluorescence change, even if there was a labeled amino acid in position to be removed. This corresponds to an apparent insertion of a non-labeled amino acid into the fluorosequence. Note that the use of a fluorescing Edman reagent (e.g., DABITC or FITC [25]) would enable direct monitoring of every coupling and decoupling step of the chemistry, providing an internal error check for successful completion of the Edman cycle as in Figure 1. Nonetheless, non-fluorescent Edman reagents such as phenylisothiocyanate are much more commonly used, and we therefore investigated the effect of this parameter.

### A framework for modeling single-molecule sequencing under non-ideal conditions

To analyze how peptide sequencing efficiency is affected by the above three types of errors and to map fluorosequences to source proteins, we developed a modeling framework to simulate the process. Unlike the ideal case where fluorosequences are faithful to their source peptides, and hence mapping to the reference proteome is trivial, accounting for errors such as the three previously highlighted complicates mapping. For example, the fluorosequence “xKxxK” cannot be uniquely attributed to the “GK*EGK*” peptide, since Edman failure at the first position of peptide “GGK*EGK*” or a fluorophore failure on the first lysine of “K*K*EGK*” could also yield the same pattern. While errors arising from the inefficiency of Edman chemistry and fluorophore failure are tractable by analytical solutions, the non-Markovian nature of photobleaching events forces us to employ a Monte Carlo approach.

We therefore developed a Monte Carlo procedure to simulate thousands of copies of each of the 20,252 proteins in the human proteome being subjected *in silico* to fluorosequencing in order to obtain a random sample of the fluorosequences produced for a specified set of error rates. Figure 4 details the simulation steps; the Methods provide more complete descriptions of the error models and pseudo-code for the overall procedure.

**Figure 4:**
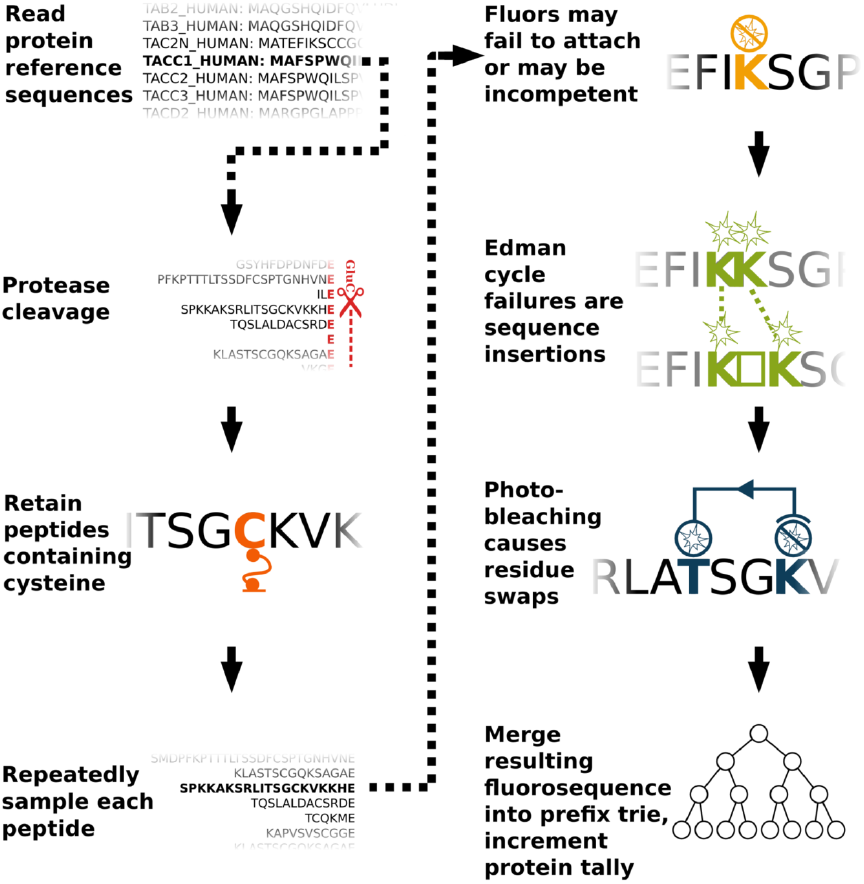
Overview of a Monte Carlo simulation of fluorosequencing with errors. In detail, (A) a protein sequences is read as amino-acid character strings from the UniProt database. For each protein sequence, the subsequent steps are repeated: (B) Proteolysis was simulated and (C) peptides lacking the residue for surface attachment (e.g. cysteine) were discarded. (D) All remaining peptides were encoded as fluorosequences and subsequent steps were repeated in accordance to the desired sampling depth: (E) The fluorosequences were altered *via* random functions modeling experimental errors - (1) labels were removed modeling failed fluorophores or failed fluorophore attachment, (2) positions of the remaining labels were randomly dilated modeling Edman reaction failures, and (3) fluorophores were shifted upstream from their positions, modeling photobleaching. (F) Each resulting fluorosequence was sorted based on its position and label type and merged into a prefix trie to tally the frequencies of observing each fluorosequence from a given source protein.

Each sample observation generated by the Monte Carlo simulation is a sequence of luminosity drops yielded by one individual peptide subjected to *in silico* Edman cycles. We conservatively assume that we cannot observe or infer the absolute number of fluorophores labeling a peptide, but that we can monitor and statistically discriminate whether, after each attempted Edman cycle, there has been a decrease in luminosity in each fluorescent channel, consistent with signals previously shown to be discernable for single molecules [17]. For the purpose of the simulation, we make the simplifying assumptions that different fluorophores have fully distinguishable signals, do not exhibit fluor-to-fluor interactions or Förster resonance energy transfer, nor exhibit channel bleed-over.

The fluorosequences (observed reads) from the simulations are next collated into a prefix trie [26], as illustrated for a simple example in Figure 5. Each fluorosequence is linked in the trie to its source protein(s) and associated count(s) of observations over the course of the simulation, thereby empirically estimating the fluorosequence’s source protein probability distribution. Figure 6 illustrates two extreme cases of protein probability distributions for a given fluorosequence. Importantly, modeling the frequency of source proteins for fluorosequences is equivalent to obtaining (within sample error) the posterior fluorosequence-to-protein probability mass functions – *i.e.* the set of probabilities *P*[*p*_*j*_|*f*_*i*_] that a given fluorosequence *f*_*i*_ was caused by source protein *p*_*j*_ (henceforth called the *attribution probability mass function (p.m.f.)*) Notably, by sidestepping problems associated with developing algorithms for inverting fluorosequences to their source peptides, and the peptides’ own derivation from source proteins, we make the strategy amenable for incorporating additional experimental parameters, including fluorophore spectral channel bleed-over or protease inefficiencies. Thus, the attribution p.m.f.’s provide a natural framework both for modeling errors and for directly mapping actual experimentally observed fluorosequences to proteins in the proteome. A fluorosequence can be associated with the protein most likely to yield it based on properties of this distribution, for example by applying a confidence threshold (see Methods).

**Figure 5:**
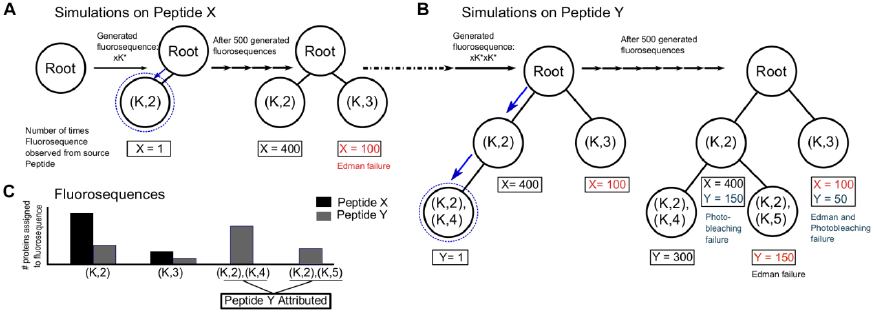
A simple example of the trie structure for storing and attributing fluorosequences to peptides or proteins. Consider a toy peptide mixture with peptide X (sequence GK*EGC, where K* represents fluorescently-labeled lysine; the sequence can be simplified to (K,2)) and peptide Y (GK*GK*EC; represented as (K,2),(K,4)). Panels (A) and (B) summarize populating the trie with fluorosequences from 500 copies each of Peptide X and Y, respectively. For example, peptide X might generate fluorosequence xK*, incorporated into the trie as a new node (K,2), indicated by the dashed blue lines and arrows in panel (A). (B) Simulations on Peptide Y add additional nodes to the trie. For example, the fluorosequence xK*xK* yields an additional node (K,2),(K,4) after traversing node (K,2). Additional fluorosequences are incorporated into the trie in a similar fashion, along with a tally of the number of observations of each fluorosequence, stored for each trie node along with the source peptide identities. Following the Monte Carlo simulation, the frequency of each source protein or peptide can be calculated for each trie node. To simplify data analysis and visualization, thresholds can be applied (see **Methods**) to identify and count those source proteins most confidently identified by the observed fluorosequences. Here, fluorosequences ((K,2),(K,5)) and ((K,2),(K,4)) confidently identify peptide Y, while Peptide X is less confidently identified by fluorosequences (K,2) or (K,3).

**Figure 6:**
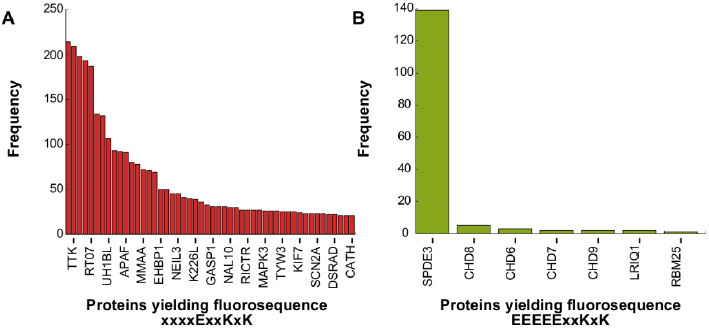
Monte Carlo sampling reveals the confidence with which fluorosequences can be attributed to specific source proteins. (A) and (B) represent two example fluorosequences, illustrating opposite extremes in terms of the number of proteins capable of yielding each sequence. In (A), the frequencies with which rival source proteins yield fluorosequence “xxxxExxKxK” in the Monte Carlo simulations indicates low confidence in attributing that fluorosequence to any one protein. In (B), a single protein is by far the most likely source of fluorosequence “EEEEExxKxK”. (X-axes represent incomplete lists of proteins, ranked ordered by the frequency they are observed to generate the given fluorosequence in the simulations.)

In future applications of attribution p.m.f.’s to interpret fluorosequencing data from real samples, one might also wish to model realistic numbers of copies per protein processed through the simulation pipeline, since the Monte-Carlo based deconvolution of fluorosequences to source proteins will be affected by protein abundance dynamic range as well as simulation depth. For example, high simulation depth would not only reduce the sampling errors, but also accurately attribute low abundance proteins from confounding high abundance proteins that generate the same fluorosequence by a low probability event. Although we did not explore this aspect, simulating protein copies based on their prior known abundances [27] might significantly reduce Monte-Carlo simulation computational resources. The version of the simulation we performed here makes no such assumptions about protein abundance, and thus corresponds to a Bayesian flat prior expectation on protein abundance, applicable to any sample.

### More amino acid colors compensate for photobleaching and poor Edman efficiency

Using the Monte Carlo scheme, we simulated sequencing the human proteome to a simulation depth of 10,000 copies per protein, performing a parametric sweep of 216 experimental parameter combinations (corresponding to six values for each of the three error parameters). Figure 7 illustrates the effects for three alternate labeling schemes of varying Edman efficiency and fluorophore half-life on the percentage of proteins identified after 30 Edman cycles, given fluorophore failure rates ranging between 0 and 25%. As in Figure 2, diversifying the labels offers the greatest improvement in proteome coverage, even with relatively poor process efficiencies.

**Figure 7:**
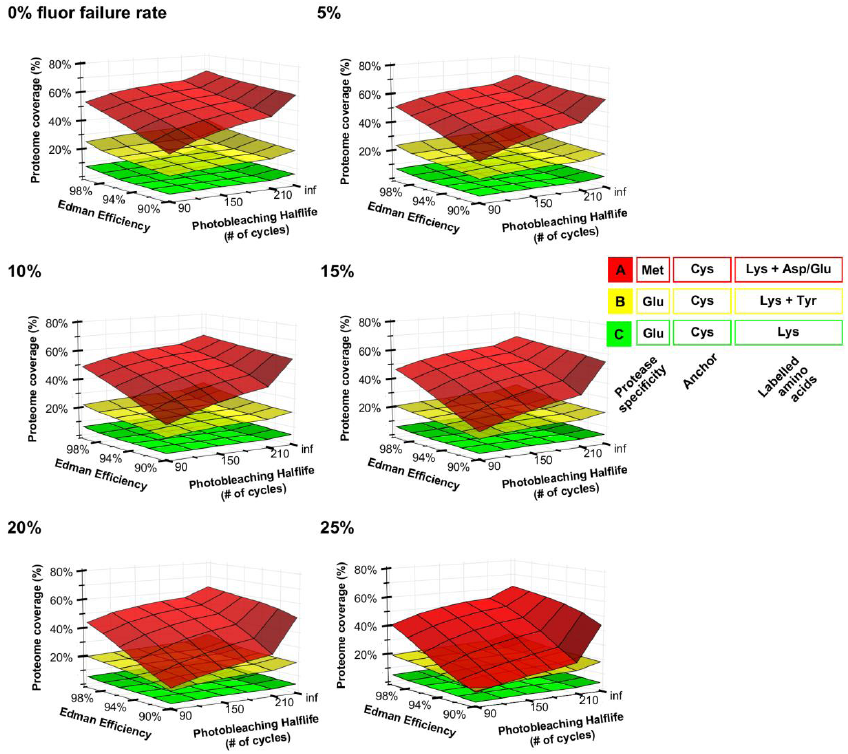
Surface plots illustrate the consequences of differing rates of Edman efficiency, photobleaching, and fluorophore failure rates. Each panel summarizes the consequences of varying rates of photobleaching and Edman failures for a different fixed fluorophore failure rate, ranging from 0% to 25%, as calculated after simulating 30 experimental cycles on the complete human proteome at a simulation depth of 10,000 copies per protein. Photobleaching shows the strongest negative impact on proteome coverage when compared to other errors; increasing the number of distinguishable labels strongly increases proteome coverage. Labeling and immobilization schemes are denoted as in Figure 2. For comparison, literature evidence suggests that common failure rates of fluorophores may be about 15-20% [18,32], Edman degradation proceeds with about 94% efficiency [33], and the mean photobleaching lifetime of a typical Atto680 dye is about 30 minutes [23], corresponding to 1800 Edman cycles, assuming 1 sec exposure per Edman cycle. Thus, we expect error rates to be sufficiently low for effective fluorosequencing.

Encouragingly, the number of proteins identified is reasonably robust to changes in fluorophore failure rates. For example, a 25% increase in failure rate causes only a 0.8%-6.4% reduction (range includes all parameter combinations) in proteome coverage for schemes A and B (see Figure 7 for scheme descriptions). However for scheme C, a 25% increase in fluorophore failure rate causes a 19% reduction in proteome coverage under moderate estimates of photobleaching and Edman efficiency. Scheme C is less robust *vis-à-vis* all simulated errors because the boost in the positional information stemming from abundant aspartates and glutamates is rapidly undermined by experimental errors, as there are higher chances for other peptides to confound the fluorosequence.

Notably, the photobleaching half-life has the greatest effect of any of the tested parameters on protein identification, causing up to 50% loss in proteome coverage (under scheme C). The steepest decrease in the number of proteins identified occurs when photobleaching is considered (from infinite cycles to 210 cycles) and tapers with lower half-life. Although photobleaching shows the strongest impact of any of the errors considered, it is worth noting that the half-lives of commercially-available fluorophores are sufficiently longer than those simulated that we anticipate that this error source will not derail a real implementation of fluorosequencing. For example, the widely used Atto680 dye has a mean photobleaching lifetime of about 30 minutes [23], corresponding to 1800 Edman cycles, assuming 1 second exposure per Edman cycle. Oxygen-scavenging systems are also widely used in single molecule imaging experiments to reduce the effects of photobleaching [19]. Thus, the most critical error rates appear to fall within acceptable ranges, supporting the feasibility of fluorosequencing.

### Determining the positional information of amino acids as a general principle for next-generation protein sequencing

The fluorosequencing method relies on the positional information of specific subsets of amino acids within peptide sequences. The scheme can be generalized as a framework fulfilling two conditions – (a) *an observable event* ‘e’, which occurs by detection of a known single amino acid or a class of amino acids, and (b) *a sequential analytical process*, which increments or decrements the sequence in a known direction and by constrained number of amino acids. For the event, while we have suggested the example of detecting fluorescently labeled amino acids, other modalities might be considered, such as detecting voltage changes or reactivity of monitored amino acids. Besides Edman degradation, other valid sequential processes could include sequential treatment with known sequence specific peptidases or directional protein translocation through a nanopore channel [9] at a defined translocation rate. The monitoring of sequenced detection events gives information-rich patterns (such as “x-e-e-x…” where ‘x’ is a non-event denoting one or more amino acids) capable of being mapped back to a reference proteome. The nature of this information lies between the extremes of information content, wherein every amino acid corresponds to a distinct event or there is no observable event with the process (as, for example, a peptide translocating through a channel but not generating a detectable signal). In principle, many event-process strategies might be suitable for peptide sequencing and interpretation using a scheme similar to the one we present.

## CONCLUSIONS

We propose a strategy for the parallel identification of proteins in a complex mixture based on the positional information of amino acids in peptides. The integration of a 60-year-old, highly optimized Edman chemistry [11] with recent advances in single-molecule microscopy [28] and stable synthetic fluorophore chemistry [29] makes this strategy particularly amenable for experimental execution in the near future. Modeling of experimental errors suggests this strategy can be reasonably expected to identify a high percentage of the proteome, comparable to mass spectrometry, and potentially brings the advantages of single molecule sensitivity and—if next-generation single molecule sequencing is a reasonable proxy—throughputs of hundreds of millions or billions of molecules sequenced per run. Monte-Carlo simulations provide a framework to accommodate the inevitable experimental errors and probabilistically identify proteins from the observed fluorescent patterns. Successful experimental execution of the proposed strategy will not only lead to progress in proteomics, but enable progress in engineering and chemistry to enable the technology.

## METHODS

### Datasets

The UniProtKB/Swiss-Prot complete *H. sapiens* proteome (manually reviewed) was downloaded on 29^th^ May 2013 and used for all simulations, comprising 20,252 protein sequences and ignoring alternatively spliced isoforms.

### Monte Carlo simulations

Simulations were programmed in Python using Merseinne Twister [28] as the source of randomness, and implemented in parallel using the Texas Advanced Computing Center. For the purposes of simulation, the proteome can be considered dictionary pairs of protein identifiers and amino acid sequences. We began the simulations with 10,000 copies of each protein sequence. The first two steps in the simulation split each amino acid sequence string at residue(s) corresponding to the protease specificity (*e.g.* E for the GluC protease) and then discard substrings that lack the anchor residue (*e.g.* substrings not containing C). Alternating Edman degradation steps and TIRF observations on the resulting peptides provide temporal ordering for luminosity drops, resulting in an observed fluorosequence for each peptide. In the simulation, fluorosequences were initialized from amino acid substrings’ correct fluorophore positions, and experimental errors were then introduced sequentially, modifying the fluorosequences in accordance with each type of error’s appropriate probability distribution.

The three experimental sources of error sources were modeled into the Monte Carlo simulation as follows:

1. Inefficient dye labeling - The probability of an amino acid not being labeled with its intended label or being labeled with a nonfunctional dye (*i.e.* a dye that attaches but is incapable of fluorescence) is modeled as a Bernoulli variable. For each label prepared for the experimental procedure, there is a probability *u* that the fluor will never be observed.
2. Edman degradation is represented as an attempt to remove one amino acid residue per cycle. These attempts are modeled as a Bernoulli process, since every experimental cycle is independent of the preceding cycle. The probability of the N-terminus amino acid being successfully cleaved off is assigned a parameter *p* and the corresponding failure follows as *q* = 1 − *p*. Failure of Edman chemistry delays the removal of a downstream labeled amino acid by one experimental cycle, and thus dilates the inter-label intervals in the fluorosequence. Using this model, the probability that an inter-label interval *d* requires *d* + *e* experimental cycles before the subsequent label is removed is 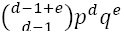. A random number is drawn from this distribution to indicate the dilation for each interval. We assume that Edman chemistry stops at the first cysteine from the N-terminus.
3. Photobleaching is the irreversible photo-induced destruction of a fluorophore. The photobleaching process can be best described as a stochastic phenomenon and modeled by an exponential decay function [30]. Every fluorophore has a defined half-life based on solvent conditions and laser operating conditions [31]. The periodic laser excitation has an additive effect on the fluorophore’s half-life: exciting a fluorophore once for thirty seconds and, after an arbitrary delay, again for a further thirty seconds will photobleach the fluorophore with the same probability as a continuous excitation for one minute. We assume a constant period of laser exposure per experimental cycle. To model whether labeled amino acids have been cleaved, the probability of a fluorophore still on the peptide surviving *k* experimental cycles can be modeled as an exponential decay *e*^-*bk*^, where *b* is an experimentally-determined characteristic constant of the fluor being used, *k* is the number of experimental cycles performed, and *e* is Euler’s constant. We shift labels to earlier experimental cycles based on random numbers drawn from this exponential decay.

For a given simulation, all simulated fluorosequences were collated into a prefix trie whose keys were the sequences of luminosity drops and associated values represented the counts of source proteins yielding those fluorosequences. One trie was calculated for each given choice of error rates, protease and labels, based upon simulating 30 Edman cycles of fluorosequencing 10,000 copies of each protein in the human proteome. For each fluorosequence in the resulting trie, its source proteins were counted, allowing proteome coverage to be calculated.

The simulation can be summarized as pseudo-code:

INITIALIZE **result trie** as an empty prefix trie.

FOR **protein** IN **proteome**:

**peptides ←** Proteolyse **protein** at the carboxyl side of a given amino acid corresponding to the protease used.
FOR **peptide** IN **peptides**:

Discard **peptide** if it does not contain at least one occurrence of the amino acid for anchoring to the surface.
FOR **peptide** IN **peptides** REPEAT **10000** TIMES:

Attach labels to amino acids with a given probability. Labeling probability is uniform and mutually independent for all amino acids.
Adjust positions of labeled amino acids to reflect possible Edman failures. All Edman reactions for each individual peptide have a uniform probability of success specified by a given parameter, and are mutually independent. We assume the Edman reaction cannot proceed past the first amino acid hybridized to the surface.
Adjust positions of labeled amino acids to reflect potential photobleaching. All fluors’ survival functions are mutually independent exponential decays characterized by a given photobleaching constant.
Collate final sequence of **tuples (fluorosequence)** for this **peptide** into the **result trie**.

Traverse the trie. For each node, find the most frequent source protein yielding that fluorosequence. For the purposes of data visualization, if the most frequent protein yielded the fluorosequence at least ten times, and all other source proteins for that fluorosequence combined are responsible for less than 10% of all observations, then that fluorosequence is considered to be uniquely attributed to the protein.

RETURN the set of proteins that have at least one uniquely attributed fluorosequence.

More detailed pseudo-code is also provided as a supplemental file.

A parameter sweep was performed for the three labeling schemes as in Figure 7 at a simulation depth of 10^4^ copies per protein, sweeping 216 experimental parameter combinations (testing six values for each of the three error parameters described) spanning fluorophore failure rates of 0%, to 25%, photobleaching half-lives from 90 minutes to infinity (*i.e.,* no photobleaching), and Edman degradation efficiencies from 90% to 100%.

### Attributing fluorosequences to peptides and proteins

For more efficient use of computer memory, trie structures were calculated separately for multiple subsets of the proteome and the resulting tries merged before analysis by traversing all fluorosequences in each trie, and adding each fluorosequence along with its protein counts into a master trie for that simulation. Then, the counts of each fluorosequence and affiliated peptide were analyzed to calculate a frequency distribution of the number of times peptides from a given source protein generated a given fluorosequence. For the purposes of summarizing the data, two criteria were applied to this distribution to attribute a fluorosequence uniquely to the protein: (a) its primary source protein yielded the fluorosequence at least 10 times out of a 10^4^ simulation depth, and (b) the summation of frequency from all other source proteins were responsible for less than 10% of that fluorosequence’s occurrences. While the former criterion addresses sample error, the latter addresses confounding from other proteins.

The core Python script and C module used for implementation of the Monte Carlo simulation are attached as supporting material.

## ACKNOWLEDGMENTS

We are deeply grateful to Andrew Emili for posing the challenge to sequence single peptide molecules, and to fruitful discussions with Andrew Ellington, Erik Hernandez, Jeff Pruet, Eric Anslyn, Brian Cannon, Rick Russell, William Press, Joseph Marotta, and Andrew Horton in the course of developing these ideas. The authors acknowledge Angel Syrett for illustrating Figure 1, and the Texas Advanced Computing Center at the University of Texas at Austin for providing high performance computing resources.

### FINANCIAL STATEMENT

J.S. is a Howard Hughes Medical Institute International Student Research fellow. This work was supported by grants from NIH, NSF, CPRIT, and the Welch foundation (F1515) to E.M.M. The funders had no role in study design, data collection and analysis, decision to publish, or preparation of the manuscript.

